# Rete Ridge Topography Regulates Epidermal Stem Cell Differentiation Programs and Chromatin Accessibility

**DOI:** 10.64898/2026.04.07.716516

**Authors:** Rui Fang, Ryoko Hamaguchi, Shuyun Xu, Justin Xu, Wonhye Lee, Kristina Todorova, Stefano Sol, Xunwei Wu, Wanqiao Gong, Mindy Nguyen, Jacob Shi, Alvaro C. Laga, Seung-Schik Yoo, George F. Murphy, Anna Mandinova, Christine G. Lian

**Author notes:** Corresponding author Christine G. Lian.

## Abstract

Stem cell niches regulate tissue homeostasis through biochemical and physical cues. Epidermal stem cells (EpiSCs) reside preferentially in the concave bases of rete ridges, which flatten with skin aging. Whether loss of this niche geometry contributes to EpiSC dysfunction in aging is unknown. We engineered flat and curved collagen matrices by 3D-bioprinting and found that concave topography promoted EpiSC differentiation and suppressed cell-cycle programs, with chromatin accessibility changes enriched for motifs of differentiation-associated transcription factors. Single-cell RNA-seq analysis of young and aged epidermis revealed shared age- and curvature-related gene signatures and changes in master differentiation regulators, supporting the physiological relevance of our ex vivo model. Immunofluorescence staining showed fewer KLF4^+^ and GRHL3^+^ granular keratinocytes and the emergence of SOX9^+^ basal cells in aged epidermis, consistent with the curved/flat model. Thus, rete ridge curvature is a key physical cue that shapes EpiSC fate, and its loss may contribute to epidermal aging.

## INTRODUCTION

Chronologic skin aging is characterized by functional impairment, including reduced epidermal keratinocyte proliferation (*1–3*), compromised barrier function *(4-6*), and clinical and histologic epidermal thinning and atrophy (*2, 7, 8*). A key histological feature of youthful skin is the presence of rete ridges, honeycomb-like downward projections ranging 50–400 μm in width and 50–200 μm in depth that interdigitate with dermal papillae to form the dermal–epidermal junction (*9–11*). With aging, this architecture becomes progressively flattened, coincident with loss of morphological complexity inherent in epidermal atrophy (*2, 7, 8*).

Epidermal stem cells (EpiSC) are slow-cycling basal keratinocyte subpopulations with self-renewal capacity and differentiation plasticity that preferentially localize within the concave bases of epidermal rete ridges (*12–14*). Within these spatially defined microenvironments, EpiSC perform essential stem cell functions required to maintain epidermal homeostasis and initiate repair, as demonstrated by morphometric and stem-cell marker molecular studies (*15–17*). In contrast, non-stem keratinocytes reside in flatter intervening regions along the dermal–epidermal junction where they persist with age as stem cell–rich rete ridges flatten and undulations characteristic of youthful skin are lost.

Despite extensive characterization of age-associated stem cell decline, it remains unclear whether intrinsic stem cell attrition drives architectural deterioration, or alternatively, whether atrophic epidermis leads to primary topographical alterations in the stem cell niche resulting in secondary stem cell dysfunction and depletion. Emerging studies underscore the role of biophysical cues—including matrix stiffness, mechanical forces, and topographic features of the extracellular environment—in regulating stem cell fate decisions (*18–20*). Indeed, mechanical regulation of cell fate in the epidermis has recently been highlighted as a central determinant of certain stem cell properties and lineage commitments (*21–23*).

Here, we show that rete ridge niche curvature is an intrinsic regulator of human epidermal stem cell fate. Using 3D bioprinting, we engineered anatomically accurate concave collagen matrices that precisely recapitulate the curved geometry of youthful rete ridges and compared them with flat substrates modeling aged epidermis. We find that concave niche topography promotes epidermal differentiation, suppresses cell-cycle programs, and induces distinct chromatin accessibility states in epidermal stem cells. These data identify a physical property as fundamental as niche curvature as an important physical cue governing epidermal stem cell behavior and suggest that loss of rete ridge architecture may directly contribute to epidermal stem cell dysfunction that typifies skin aging.

## RESULTS

### 3D-bioprinted geometric models of EpiSC niche in young and aged human skin

One of the most striking and consistent histological features observed in aged human skin is marked flattening and atrophy of the rete ridge structures compared to the well-defined undulating rete ridges seen in young skin samples and the reduction of CK15^+^ epidermal stem cells at the epidermal basal layer that have been well documented in the literature (*2, 7, 24*) (representative in **Figure 1A/B**). Examining normal skin samples of young (< 30 years old, n=6) and elderly individuals (> 75 years old, n=6) (**Table S1**), we observed that epidermal thickness measured at rete ridges was reduced by more than half in aged skin (125.2±4.3 vs. 51.7±3.1 μm; P < 0.001), and was ~40% thinner in inter-ridge regions (58.3±3.1 vs. 37±6.7 μm; P < 0.05) (**Figure S1**). We sought to determine whether rete ridge geometry is a determinant of epidermal stem cell maintenance and function, and whether its loss contributes to the differences between young and aged human skin.

**Fig 1.**
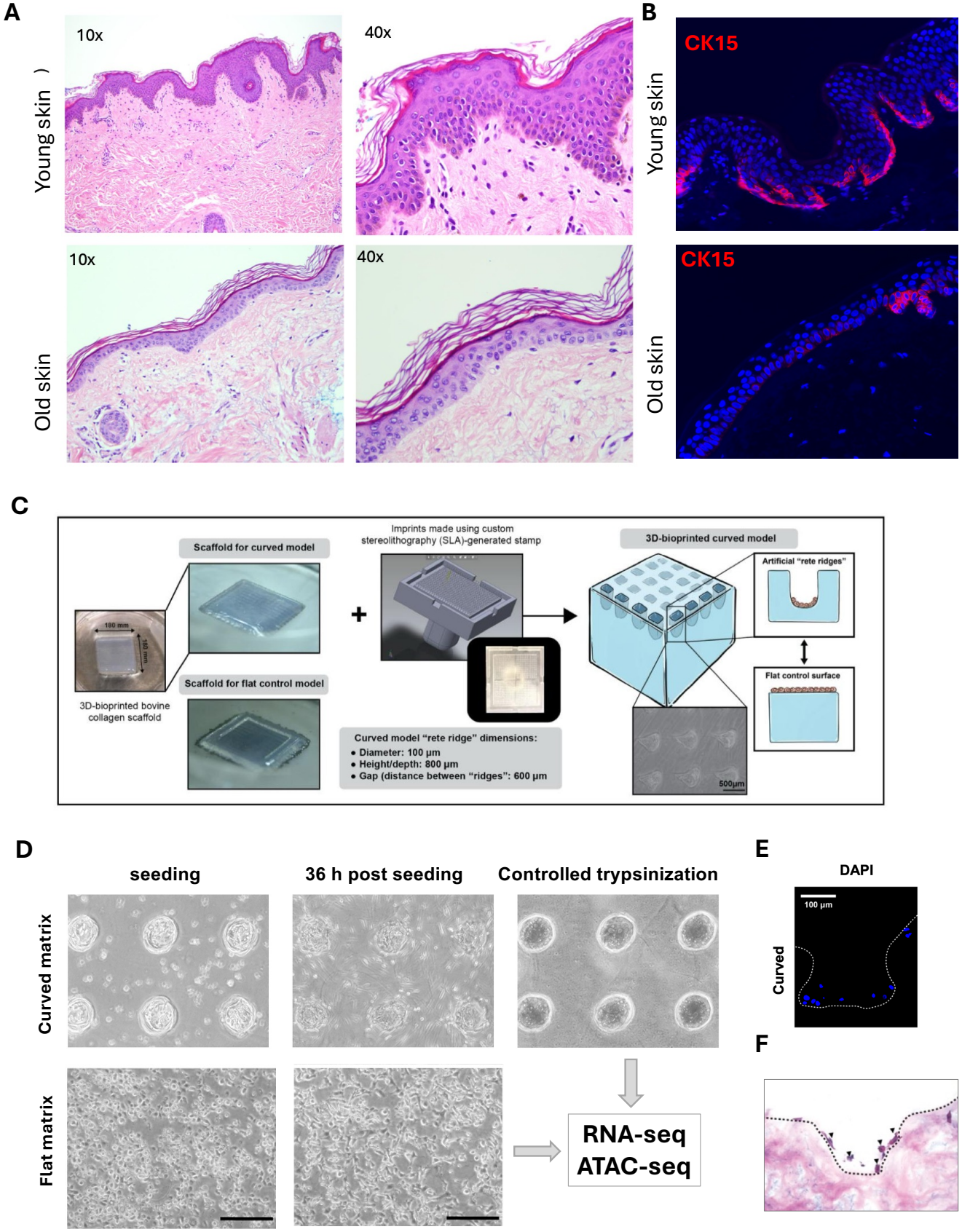
Age-associated flattening of rete ridges and construction of a 3D-bioprinted rete ridge microenvironment modeling the epidermal stem cell niche. **(A)** Representative H&E staining of young (19 years) and aged (92 years) human skin at 10× and 40× magnification. Young skin shows prominent rete ridges with deep epidermal invaginations, whereas aged skin shows epidermal atrophy with a flattened dermal–epidermal junction. **(B)** Immunofluorescence staining of CK15 in young and aged human skin. **(C)** Schematic of the fabrication of the engineered rete ridge microenvironment. A collagen scaffold was 3D-bioprinted and imprinted with a custom stereolithography-printed resin stamp to create a curved topography mimicking the rete ridges. Flat scaffolds served as controls. **(D)** Workflow for cell seeding and selective harvesting on curved and flat matrices. Shown are bright-field images of EpiSCs 36 h after seeding on flat and curved matrices, and en face views of cells in the concavities after cells on the flat surface were removed by controlled trypsinization. **(E-F)** DAPI counterstaining (E) and H&E staining (F) of cryosections transecting the artificial “rete ridge” niches. Arrows mark cells in the curved niche.

To model the geometric features of the human epidermal stem cell niche, we fabricated collagen matrices incorporating anatomically scaled concavities that reproduce the normal architecture of rete ridges in young skin. A resin stamp was designed and engineered using stereolithography 3D printing to imprint collagen scaffolds with anatomically accurate rete ridge– like topography. The stamp (18 mm × 18 mm) featured protrusions with a diameter, height, and spacing of 100 μm, 600 μm, and 800 μm, respectively, mimicking dimensions observed in native healthy human rete ridges (**Figure 1C**). Multilayered collagen scaffolds were printed on 60-mm cell culture dishes using a layer-by-layer fabrication approach to serve as the primary scaffold material for imprinting, while also providing a flat gel matrix as a control.

To determine the effect of niche architecture as a factor influencing epidermal stem cell behavior, primary human epidermal stem cells (EpiSCs) isolated from neonatal foreskin specimens were seeded onto these matrices. EpiSCs were isolated using a one-step, serum-free method described in detail previously (*25*), which were >80% positive for CK15 and <5% FABP5 positive (**Figure S2**), indicating high-quality proliferative EpiSCs with minimal contamination of differentiating keratinocytes. EpiSCs readily adhered to the gel matrix. At 36 h post-seeding, cells within the curved niche were selectively harvested by a controlled, two-step trypsinization that removed cells adhering to the surrounding flat surfaces (**Figure 1D**). A total of 100,000 cells were used to seed each gel matrix, a condition that we had optimized to avoid overseeding and minimize the formation of cell clumps in the curved niche. Immunofluorescence and H&E staining of cryosections cut perpendicular to the scaffold surface, transecting the artificial “rete ridge” niches, confirmed that cells spread mostly as a single layer within the curved niche under this condition (**Figure 1E/F**). We performed ultra-low input RNA-seq and ATAC-seq on EpiSCs harvested from the curved niche and flat matrix to determine the effects of curved and flat geometry on EpiSC transcriptome and epigenome.

### The curved “rete ridge” niche promotes epidermal stem cell differentiation

RNA-seq analysis of EpiSCs harvested from curved and flat gel matrices (n = 3 per condition) identified 1,242 up- and 1,034 downregulated genes (fold change ≥ 1.5 and FDR < 0.05), demonstrating that niche curvature had a pronounced effect on EpiSC gene expression (**Figure 2A**). These differentially expressed genes (DEGs) were enriched in cell cycle KEGG pathway, and pathways related to skin development, including cornified envelope formation, integrin signaling and ECM–receptor interaction (**Figure 2B**). Strikingly, the upregulated genes were enriched in Gene Ontology Biological Process (GOBP) pathways of epidermal development, keratinization, keratinocyte differentiation, skin barrier, intermediate filament organization, wound healing, skin development, and cell–cell adhesion (top 10 in **Figure 2C;** KEGG in **Figure S3A**). In contrast, the downregulated genes were enriched in cell division, cell cycle, DNA replication, and other pathways related to cell proliferation (top 10 in **Figure 2D**; KEGG pathways in **Figure S3B)**. Gene Set Enrichment Analysis (GSEA) confirmed the marked downregulation of the cell cycle pathway and upregulation of epidermis development pathway (**Figure 2E/F**, top 10 most up- and down-regulated pathways in **Figure S3C**). Key cell cycle genes such as *CDK1, CDK2, CCND1, CCNB2* and *AURKB* were significantly downregulated, whereas negative regulators of cell proliferation, including cyclin-dependent kinase inhibitor 2D (*CDKN2D*) and the tumor suppressor *TP73*, were upregulated (**Figure 2G**), suggesting suppression of cell division. Most DEGs in epidermis development and keratinization pathways were upregulated, including many cytokeratin genes, granular markers (*FLG, CALML5*) and terminal differentiation genes involved in cornified envelope formation (*IVL, EVPL, PPL, SCEL, CNFN, SPRR*s, *LCE*s, *TGM5* and *CASP14*) (**Figure 2H**). Several collagen and laminin genes were downregulated, and several tight junction components (e.g., *CGN, CRB3* and *TJP3*) were upregulated, indicating remodeling of cell-cell and cell-matrix interaction (**Figure 2I**).

**Fig 2.**
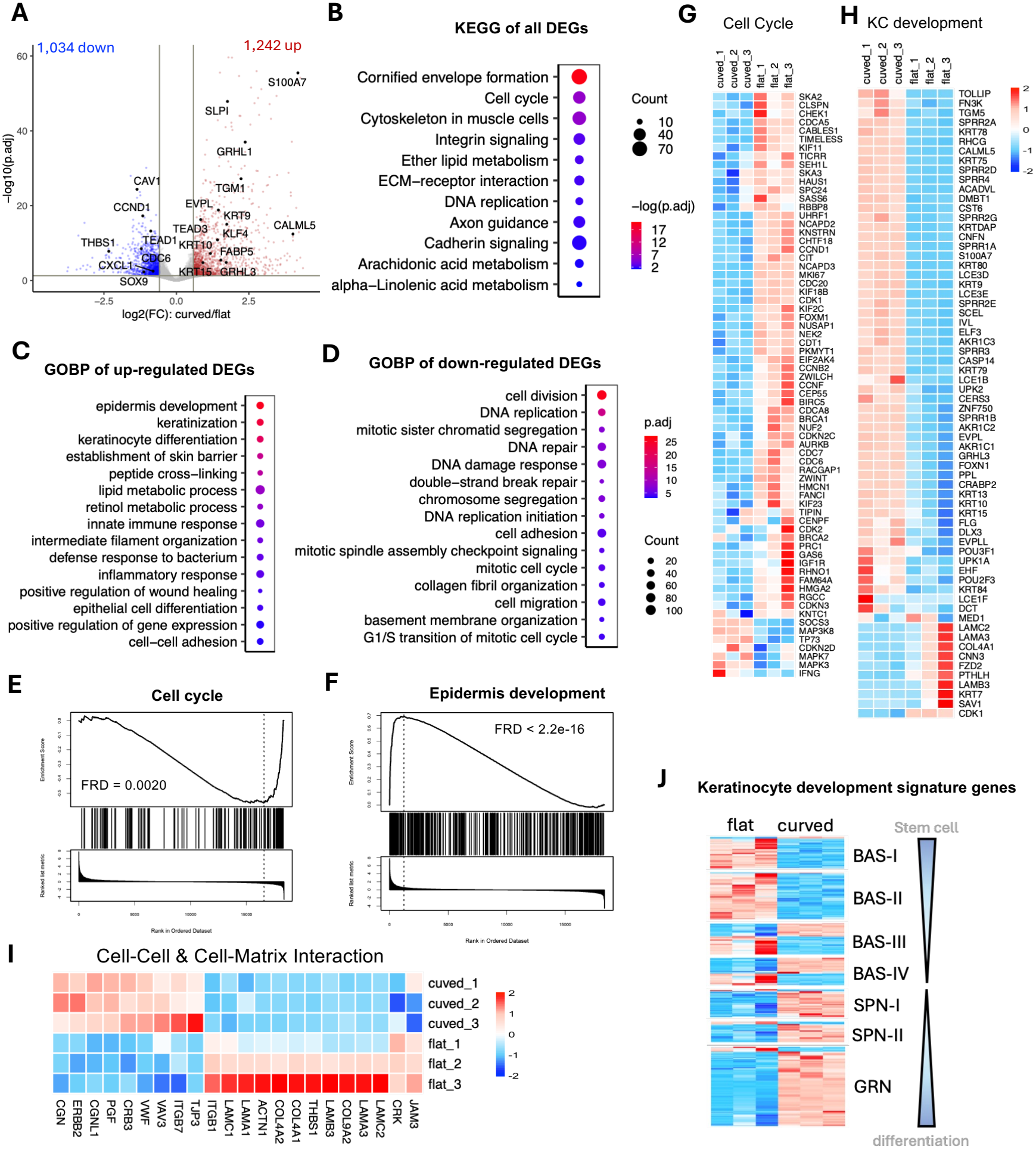
The curved “rete ridge” niche induces transcriptional programs resembling differentiating keratinocytes. **(A)** Volcano plot of DEGs in EpiSCs in the curved niche compared with those on a flat matrix. **(B)** Top 10 enriched KEGG pathways. Enrichment of up- and downregulated DEGs is shown separately in Fig S3. **(C-D)** GOBP enrichment of upregulated (C) and downregulated (D) DEGs. **(E-F)** GSEA enrichment plots of the GOBP pathways cell cycle (E) and epidermis development (F). **(G-I)** Heatmaps of DEGs in key pathways: cell cycle regulation (G), epidermis development and keratinization (H), and cell–cell and cell–matrix interaction (I). **(J)** Heatmap of expression changes of signature genes for stages of epidermal lineage progression and differentiation^26^. BAS-I to BAS-IV, basal stem cell states I– IV; SPN-I and SPN-II, spinous keratinocyte states I and II; GRN, granular keratinocytes.

These observations suggest that the curved niche has a profound effect on EpiSC transcriptome, suppressing cell replication and promoting keratinocyte differentiation. To gain further insight, we examined gene signatures identified by single-cell RNA-seq analysis of developing human EpiSC by Wang et al (*26*). They identified four distinct populations of basal keratinocyte stem cells (BAS-I to BAS-IV) in human interfollicular epidermis, and proposed a linear hierarchical model of keratinocyte differentiation in which BAS-I and BAS-II represent more stable, steady-state stem cell populations, while BAS-III and BAS-IV function as transitional states that can differentiate into spinous cells (*26*). Examining gene signatures of each keratinocyte cell stage, we observed that most of the marker genes for BAS-I and BAS-II were downregulated in EpiSCs in the curved cell niche compared to flat matrix (**Figure 2J**, top panels). Expression changes for marker genes of BAS-III and BAS-IV, the transient states of keratinocyte stem cells, were a mix of up- and downregulation and did not show a clear trend of change. Strikingly, nearly all the marker genes for spinous and granular keratinocytes, representing the committed differentiation states, were upregulated in EpiSCs in the curved cell niche (**Figure 2J**, bottom panels), consistent with the upregulation of epidermal development and keratinization pathways (**Figure 2C/H and S3**). Thus, we conclude that the curved niche preferentially promotes keratinocyte stem cell differentiation.

### ATAC-seq identifies transcription factors associated with curvature-induced EpiSC differentiation

To gain mechanistic insight into how the curved, concave niche may regulate epidermal stem cell function, we performed ATAC-seq on biological replicates of EpiSCs seeded on curved and flat gel matrices, and in conventional 2D culture (n=2 per group). ATAC-seq (Assay for Transposase-Accessible Chromatin with sequencing) is a rapid, high-resolution method for genome-wide mapping of accessible DNA, allowing the identification of active cis-regulatory elements such as promoters, enhancers, and transcription factor binding sites (*27*). A representative example of ATAC-seq signal enrichment at gene promoters and enhancers is shown at the *KRT2* locus (**Figure 3A**), which encodes keratin 2, a late differentiation keratin expressed in the upper spinous and granular layers, and was upregulated in EpiSCs in the curved niche (**Figure 3B**). The differential ATAC site at the intronic region of *KRT2* gene locus had increased ATAC signals in EpiSCs in the curved niche (**Figure 3A**, highlighted), coincided with a peak in the ReMap density track that marks consensus transcription factor (TF) binding sites derived from integrative analysis of large collections of ChIP-seq and ChIP-exo datasets (*28*). This intronic site was also marked by high H3K4me1 and H3K4me3 signals. These features are consistent with a function as an enhancer that may be important for *KRT2* gene transcription.

**Fig 3.**
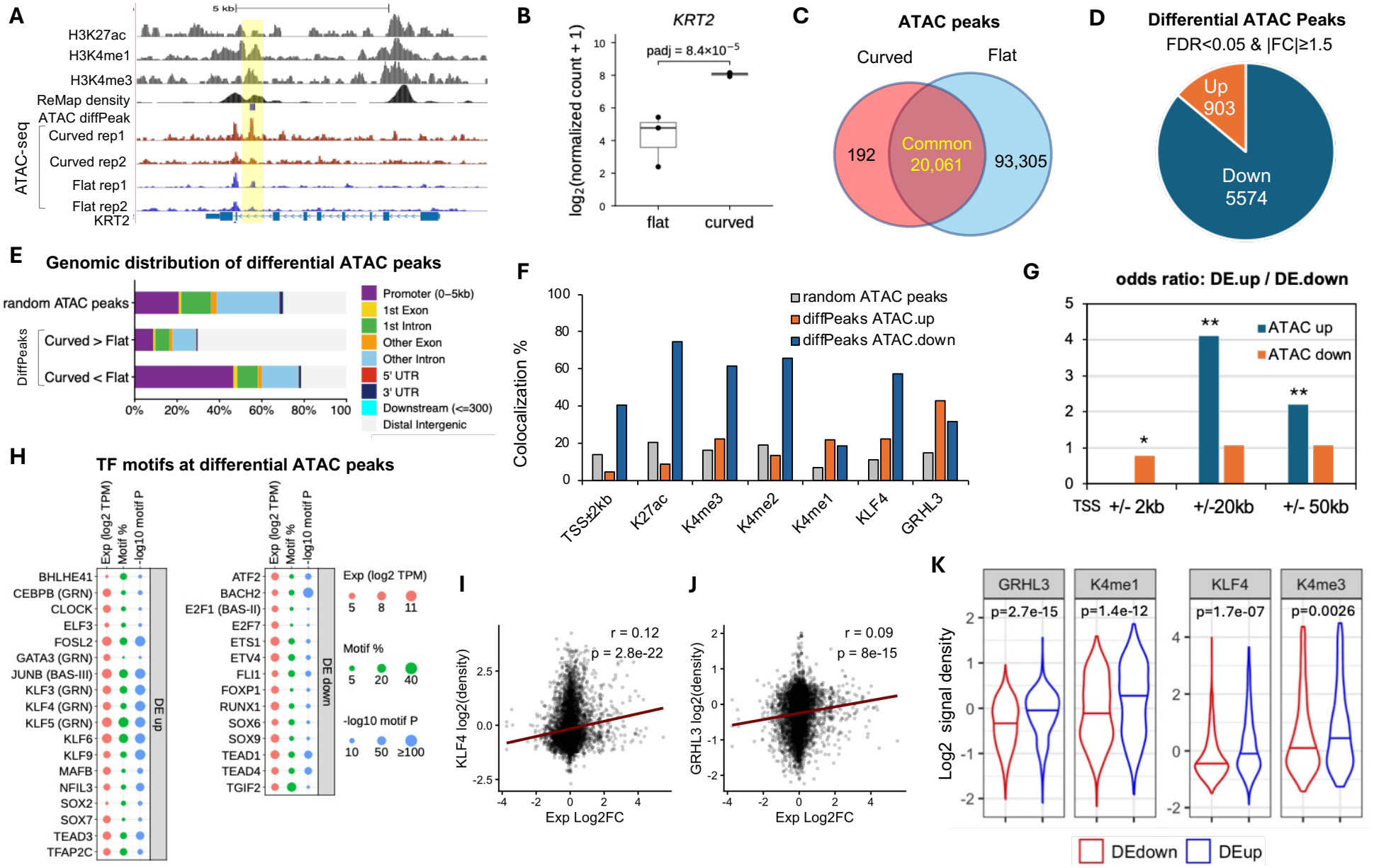
Curved niche topography induces chromatin accessibility changes that correlate with transcriptional changes. **(A)** ATAC-seq signals at *KRT2* in EpiSCs in the curved niche or on a flat matrix. ReMap, density of consensus transcription factor (TF) binding sites compiled from ChIP-seq and ChIP-exo data. **(B)** *KRT2* expression (log TPM) in EpiSCs on curved and flat matrices (n = 3 per group). **(C)** Venn diagram of ATAC peaks in EpiSCs on curved and flat matrices. **(D)** Venn diagram of differential ATAC peaks with significant increased (up) or decreased (down) ATAC signals in EpiSCs in the curved niche. **(E)** Genomic distribution of diffPeaks and of randomly sampled ATAC peaks. **(F)** Colocalization of diffPeaks with histone modifications and KLF4 and GRHL3 binding sites. Percentage of up and down diffPeaks were shown. **(G)** Preferential association of diffPeaks with DEGs. Odds ratio, ratio of up- to downregulated DEGs associated with diffPeaks, relative to all DEGs. **(H)** Motif enrichment at diffPeaks. BAS-I to BAS-IV, SPN-I and SPN-II, and GRN as in Figure 2J. DE up and down, up- and down-regulation in curved niche (vs flat; FDR<0.05), respectively. **(I-J)** Correlation between expression changes (log2FC) of the nearest genes and KLF4 (I) or GRHL3 (J) ChIP-seq signal at up diffPeaks. Pearson’s r and P values are shown. **(K)** Violin plots of KLF4, GRHL3 and H3K4me1 signals at diffPeaks associated with up- and downregulated DEGs. Pairwise t-tests on log2-transformed signals with Bonferroni correction.

Genome-wide, we identified 20,253 ATAC peaks in EpiSCs grown on curved niches using MACS2 (p < 0.001, all peaks with FDR < 0.05), nearly all of which overlapped with the ATAC peaks in EpiSCs grown on flat matrices and in conventional 2D culture (**Figure 3C & S4A**). Principal component plot showed clear separation of the 3 treatment groups (**Figure S4B**). We interrogated 403,210 loci associated with ATAC peaks in any of the 3 treatment groups using edgeR (*29, 30*) and identified 6,477 ATAC peaks with significant changes (denoted ATAC diffPeaks; FDR < 0.05 and fold change |FC| ≥ 1.5), with 903 and 5574 sites showing increased and decreased ATAC signals, respectively, in curved niche compared to flat matrices (**Figure 3D & S4C/D**). DiffPeaks with increased ATAC signals were enriched at gene promoters (46.5%), which were significantly higher compared with 20.8% of random sampling of all queried ATAC peaks (χ^2^ P < 0.001), while sites with increased ATAC signals were depleted at gene promoters (9.5%, χ^2^ P < 0.01) (**Figure 3E**). Examining histone modifications at these sites, we observed that >60% of ATAC-down diffPeaks colocalized with H3K27ac and H3K4me3, compared to 18.6% with H3K4me1, consistent with their enrichment at gene promoters (**Figure 3F**). In contrast, ATAC-up diffPeaks were depleted of H3K27ac and were enriched for H3K4me1 compared to randomly sampled ATAC peaks, suggesting a main function at enhancers (**Figure 3F**).

ATAC-seq detects open chromatin conformation, which correlates with gene activity. An increase in ATAC-seq signal intensity suggests open chromatin structure and increased chromatin accessibility for transcription factor binding; conversely, reduced ATAC signal suggests a closed chromatin conformation, which may associate with gene repression (*27, 31, 32*). ATAC-down diffPeaks that were enriched at gene promoters showed a modest preferential association with downregulated DEGs (**Figure 3G**; OR = 0.77, χ^2^ P < 0.05, odds ratio of up- to downregulated DEGs relative to all DEGs). In contrast, ATAC-up diffPeaks, which were located mostly at intergenic regions, associated with few DEGs within 2 kb of the TSS but showed a clear preference for upregulated genes at longer distances, with ORs of 4.11 and 2.19 for regions within 20 kb and 50 kb of the TSS, respectively (**Figure 3G**; χ^2^ P < 0.01 for both), consistent with distal enhancer function.

To identify transcription factors (TFs) potentially important for gene transcription and epigenetic changes in EpiSCs in the curved niche, we next performed HOMER motif analysis and identified 158 enriched motifs (p.adj < 0.001). Many of these transcription factors (TFs) are important for cell fate commitment (marked in red in **Figure S5**). Notably, KLF3, KLF4 and KLF5 of the Krüppel-like factor (KLF) family, which play important roles in epidermal development (*33–35*) and are signature genes of granular keratinocytes (GRN) defined by Wang et al. (*26*), were upregulated in EpiSCs in the curved niche (**Figure 3H**, left panel). In contrast, TFs important for EpiSC maintenance and stemness, such as SOX9 (*36*), FOXP1 (*37*) and TEAD1 (*38*), were downregulated (**Figure 3H**, right panel).

Also of interest are the Grainyhead-like transcription factor 3 (GRHL3). While not included in the HOMER motif database, *GRHL3* was significantly upregulated and expressed at high levels in EpiSCs in the curved niche. KLF4 and GRHL3 were both enriched at differential ATAC sites. KLF4 binding sites colocalized with 57.3% of ATAC-down and 22.4% of ATAC-up diffPeaks, respectively, significantly higher than 11.0% of randomly sampled ATAC peaks (**Figure 3F**). GRHL3 binding sites colocalized with 31.5% of ATAC-down and 42.7% of ATAC-up diffPeaks, significantly higher than 14.6% of randomly sampled ATAC peaks (**Figure 3F**).

To gain insight into how TFs and epigenetic marks contribute to gene regulation, we extended the analysis to a less stringent set of diffPeaks (nominal P < 0.05, |FC| ≥ 1.5; n = 12,440), which showed a similar genomic distribution pattern and preferential association with upregulated DEGs (**Figure S6**), suggesting that more modest changes in chromatin accessibility may also contribute to gene regulation. We observed a moderate but highly significant correlation of KLF4 and GRHL3 signals with expression changes of the closest associated genes, with a Pearson’s correlation coefficient of 0.12 (p<2×10^-16^) and 0.09 (p=7.99×10^-15^), respectively (**Figure 3I/J**). The signals of KLF4, GRHL3, H3K4me1 and H3K4me3 signals were significantly higher at upregulated genes than at downregulated genes (**Figure 3G**).

Taken together, these findings suggest that geometric cues influence EpiSC fate through topography-sensitive changes in chromatin accessibility and transcription factor networks, with KLF4 and GRHL3 likely playing a key role.

### Single-cell RNAseq reveals common features in young/old skin and curved/flat model

To test the physiological relevance of our model, we compared the transcriptional changes induced by curved versus flat matrices with those between young and aged human keratinocytes. We analyzed a published single-cell RNA-seq dataset of epidermis from the upper eyelid skin of young (20–30 years old, n = 3) and older (60–80 years old, n = 3) individuals (GSE281449) (*39*). Subclustering of interfollicular keratinocytes identified six populations, annotated by canonical markers as basal, basal KRT15-high, basal dividing, early spinous, upper spinous and granular keratinocytes (**Figure 4A & S7A**). Feature plots of well-established keratinocyte markers confirmed that the clusters accurately reflect distinct stages of differentiation (**Figure 4B & S7B**). KRT15 (encoding CK15) was most highly expressed in the basal and KRT15^high^ basal clusters, and was also detected in the dividing basal and early spinous clusters. In contrast, KRT2 and IVL, markers of terminally differentiated keratinocytes, were enriched in the granular cluster, as expected. GRHL3 was selectively enriched in upper spinous and granular keratinocytes (**Figure 4B & S7C**). KLF4 was widely expressed in both basal EpiSCs and differentiated keratinocytes, with the highest level in granular keratinocytes (**Figure 4B & S7C**). Importantly, the population of KLF4^+^ and GRHL3^+^ granular keratinocytes were both significantly decreased in old skin (**Figure 4C**).

**Figure 4.**
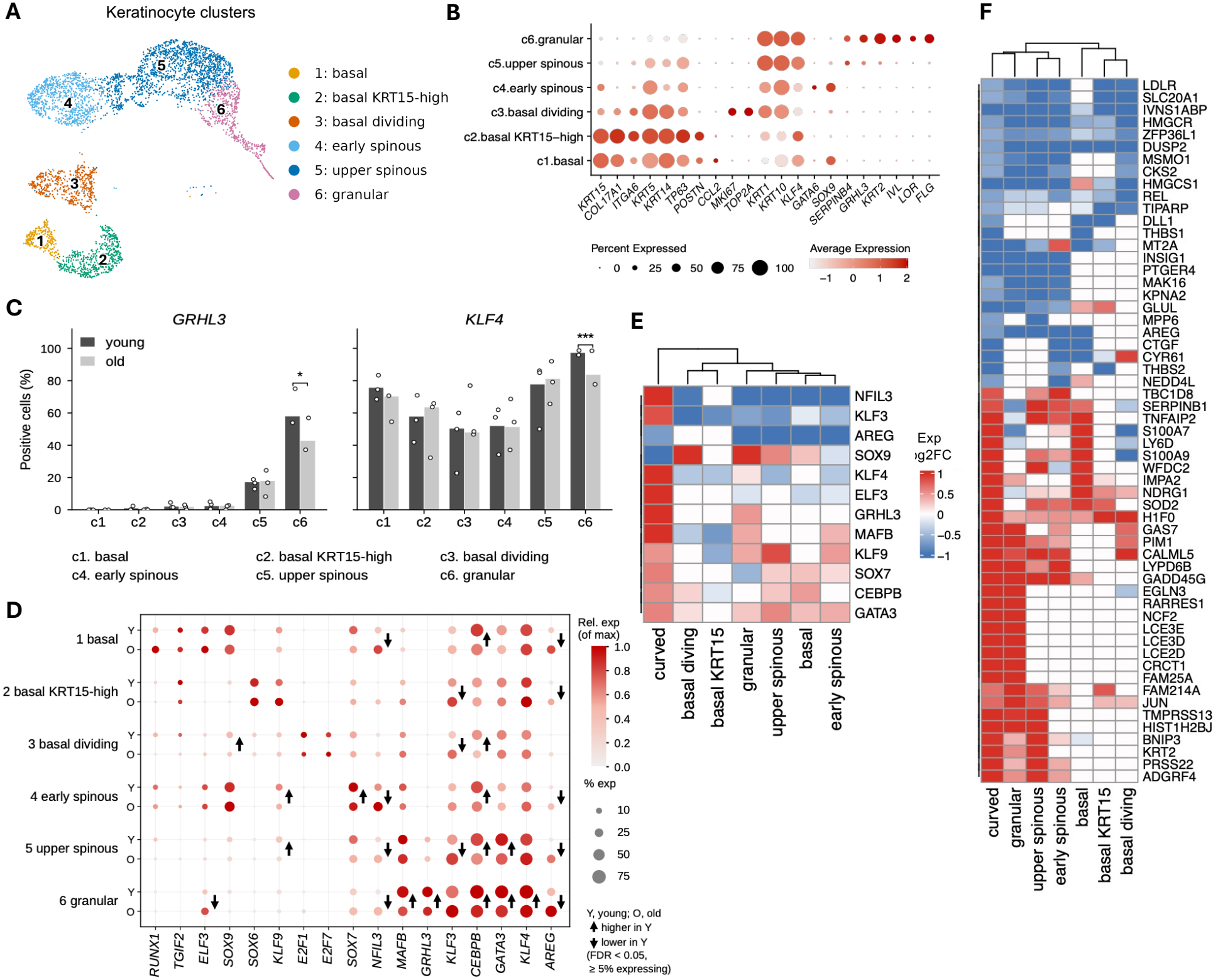
Concordant transcriptional changes in EpiSCs on curved vs. flat matrices and in young vs. aged epidermal keratinocytes. **(A)** UMAP of interfollicular keratinocyte subclusters in young and aged human epidermis (GSE281449): c1, basal; c2, basal KRT15-high; c3, basal dividing; c4, early spinous; c5, upper spinous; c6, granular. **(B)** Dot plot of signature genes of each keratinocyte subcluster. Dot size indicates the percentage of expressing cells; color indicates scaled average expression. **(C)** Population of KLF4^+^ cells and GRHL3^+^ cells in granular keratinocyte subcluster was significantly higher in young skin epidermis compared to aged skin. *, FDR < 0.05; ***, FDR < 0.001. **(D)** Dot plot of TF expression in young (Y) and old (O) keratinocytes within each keratinocyte subcluster. Arrows mark significant changes in the percentage of expressing cells (FDR < 0.05). **(E)** Heatmap of TF expression changes in young (vs. aged) keratinocytes and in EpiSCs in the curved niche (vs. flat matrix). Log2 fold changes are capped at ±1 (2-fold); changes < 10% are set to 0. **(F)** Heatmap of DEGs with concordant changes in young (vs. aged) keratinocytes and in EpiSCs in the curved niche (vs. flat matrix), with unsupervised clustering.

Examining TFs in **Figure 3H** that were differentially expressed and with motifs enriched at ATAC diffPeaks in EpiSCs in curved and flat matrices, we identified several TFs shown cluster-specific expression in human keratinocytes (**Figure 4D & S7C**). *CEBPB, GATA3, MAFB* and *KLF3* were most highly expressed in upper spinous and granular keratinocytes (**Figure 4D & S7C**). CEBPB^+^ cells were more abundant in young than in aged skin across multiple basal and differentiating keratinocyte clusters, GATA3^+^ cells were more abundant in young upper spinous and granular keratinocytes, and MAFB^+^ cells were more abundant in young granular keratinocytes (**Figure S7D**, FDR<0.05, marked by arrows). *CEBPB, GATA3, MAFB* were upregulated in EpiSCs in the curved niche and showed similar trend of changes in young skin keratinocytes (**Figure 4E**). Thus, their changes between young and aged skin are consistent with our geometric model. In contrast, KLF3 positive cell population and average expression were increased in old skin, but was upregulated in curved niche. In consistency was also observed for *NFIL3. NFIL3* was upregulated in EpiSCs in the curved niche but was more abundant in old skin, in basal and differentiating keratinocytes. *KLF9* and *SOX7* were most highly expressed spinous KCs, and the population and gene expression were higher in young skin spinous KCs, consistent with the upregulation in curved niche. SOX9 was most highly expressed in basal and early spinous KCs, and the proportion of SOX9^+^ cells did not differ significantly between young and aged skin in any KC subcluster (**Figure 4D**), but SOX9 expression trended upward in the basal dividing and granular subclusters of young skin, in contrast to the downregulation we observed in the curved niche (**Figure 4E**).

We next compared expression changes between young and aged skin with those in EpiSCs on curved versus flat matrices (**Figure 4F**). Differential expression analysis of pseudobulk counts with DESeq2 identified approximately 40–200 DEGs in each keratinocyte cluster, for a total of 199 up- and 292 downregulated genes in young versus aged skin (nominal P < 0.05 and |FC| ≥ 1.5; few genes reached FDR < 0.05). We identified 57 concordant DEGs (32 genes were upregulated and 25 downregulated) in EpiSCs in the curved niche (versus flat matrix) and in any of the keratinocyte clusters of young skin (versus aged skin). The concordantly upregulated genes included the late cornified envelope genes *LCE2D, LCE3D* and *LCE3E*, the upper spinous/granular keratin *KRT2*, the granular-layer marker *CALML5*, and the epidermal differentiation genes *CRCT1* and *FAM25A*, suggesting active terminal differentiation toward mature spinous and granular keratinocytes (**Figure 4F**). They also included the barrier-associated proteases and protease inhibitors *PRSS22, TMPRSS13, SERPINB1* and *WFDC2*, which maintain the protease–antiprotease balance required for stratum corneum maturation, desquamation and barrier integrity.

The 25 concordantly downregulated genes included genes involved in cholesterol biosynthesis and uptake (*HMGCR, HMGCS1, MSMO1, LDLR* and *INSIG1*) and the YAP–TEAD targets *CTGF* and *CYR61*. Interestingly, *AREG*, a major autocrine keratinocyte mitogen that acts through EGFR and is overexpressed in hyperproliferative skin diseases (*40, 41*), was downregulated in the curved niche and in multiple keratinocyte clusters of young skin (with a trend of down regulation in granular keratinocytes: log2FC = −1.27, P = 0.058; significant in other clusters). The proportion of AREG-expressing cells was significantly lower in young skin than in aged skin across all keratinocyte clusters (**Figure 4D & S6D**). AREG (amphiregulin) is a ligand of the epidermal growth factor receptor (EGFR) and a major autocrine keratinocyte mitogen that promotes cell proliferation in wound healing and is upregulated in hyperproliferative skin diseases (*40, 41*). Its downregulation in the curved gel matrix is consistent with the suppressed cell proliferation suggested by the transcriptomic changes we observed.

There were 75 discordant DEGs that changed in opposite directions in the two systems. Among the 64 genes that were higher in aged skin and induced in the curved niche, the stress keratin *KRT16*, the SPRR family of cornified envelope precursors (*SPRR1A/B* and *SPRR2A/D/G*), and the antimicrobial protease inhibitors *PI3* and *SLPI* are hallmarks of activated keratinocytes, which arise in response to wounding, barrier disruption or inflammation. Further supporting a stress response, aged keratinocytes showed higher expression of genes associated with DNA damage and growth arrest (*BTG2, RNF168*), the integrated stress response (*CHAC1, MTHFD2, SLC7A5*), oxidative stress (*MGST1, MT2A, MAFF/MAFK*), and inflammatory and interferon signaling (*IL18, CXCL14, CCL20, TNFAIP3, SOCS3, IFI27, GBP1*), consistent with inflammation-associated stress in aged epidermis. These differences suggest a major difference in stress response between the ex vivo geometric model and in vivo features.

In addition, the cell-surface and membrane-associated genes *ADGRF4, TNFAIP2* and *LYPD6B* were concordantly upregulated. Other adhesion-related genes changed discordantly. The basal cell–matrix adhesion genes *ITGA2* and *ACTN1* were higher in young skin but reduced in the curved niche, whereas genes associated with migratory or wound-edge keratinocytes (*LYPD3, LGALS7B* and *KANK1*) and the desmosome gene *PKP1* were higher in aged skin and induced in the curved niche. These changes suggest that cell–cell and cell–matrix interactions differ between young and aged skin keratinocytes and are remodeled in the curved niche.

### “Rete Ridge” topography-associated signatures are altered in aged human skin

Transcriptome analysis revealed significant similarities in curvature-responsive TFs and transcriptional signatures and young versus old keratinocytes, demonstrating the physiological relevance of the topography-dependent transcriptional programs identified in our *ex vivo* model. We are particularly interested in KLF4 and GRHL3, the key TFs promoting epidermal differentiation and barrier formation (*34, 42, 43*) and SOX9, which linked to stem and progenitor cell states (*36*). All these factors were differentially expressed in the curved/flat matrices (**Figure 5A**) and enriched at differential ATAC sites. We therefore examined their expression in young and aged human skin by multiplex immunofluorescence staining, co-staining with CK15 to mark basal-layer EpiSCs localized at rete ridges.

**Fig 5.**
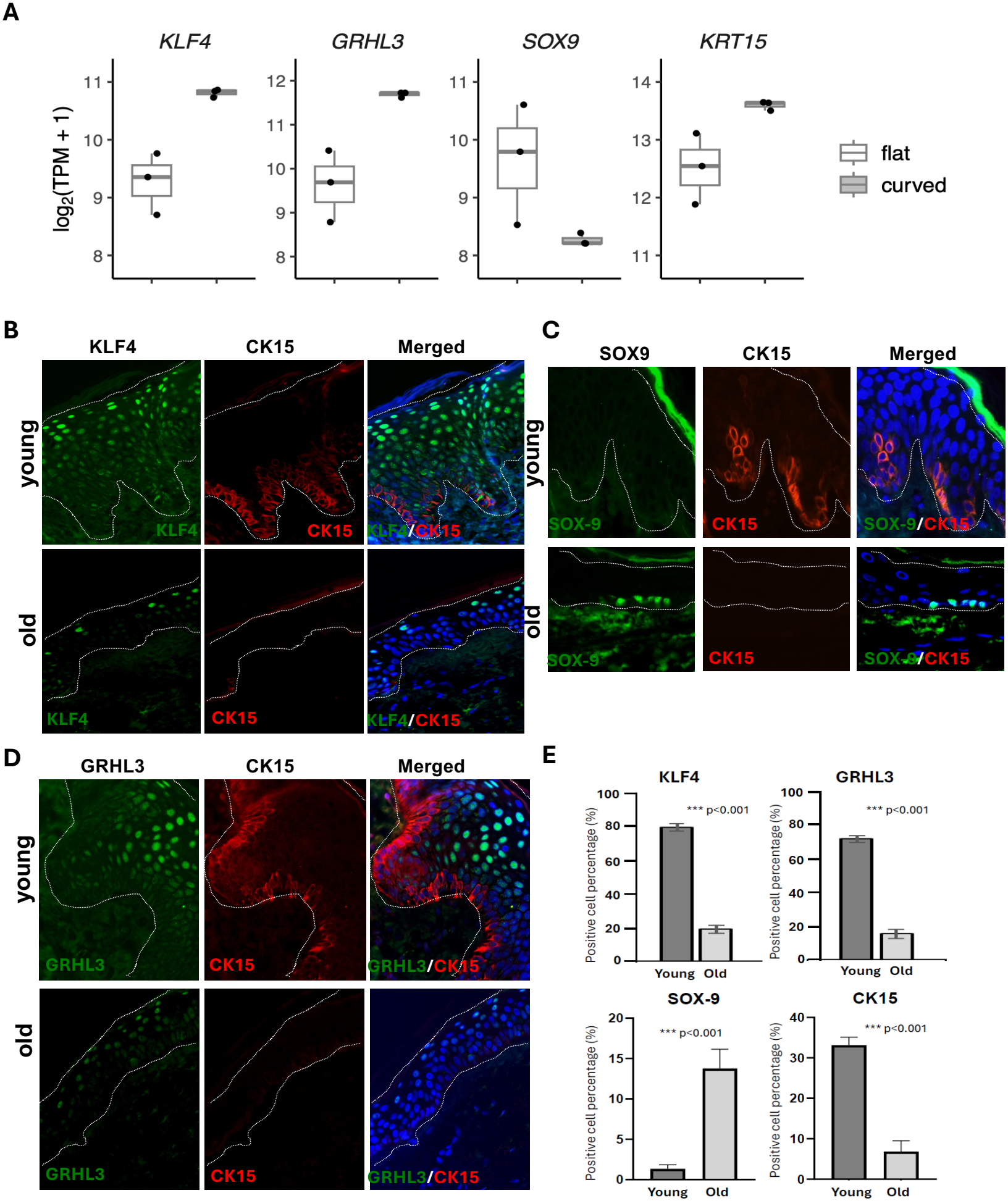
Age-associated changes in TF expression in young and aged human epidermis. **(A)** Expression of *KLF4, GRHL3, SOX9* and *KRT15* in EpiSCs on flat and curved matrices by RNA-seq (log2[TPM + 1]; n = 3 per condition). **(B)** Immunofluorescence staining of KLF4 (green) and CK15 (red) in young and aged human skin. In young epidermis, KLF4 is detected in most keratinocytes, with the highest levels in the granular layer and weak expression in CK15^+^ basal cells of rete ridges. In aged epidermis, KLF4 is reduced, particularly in basal and spinous layers, and CK15^+^ cells are diminished. **(C)** Immunofluorescence staining of GRHL3 (green) and CK15 (red). GRHL3 is highest in the uppermost layers, and the GRHL3^high population is reduced in aged epidermis. **(D)** Immunofluorescence staining of SOX9 (green) and CK15 (red). SOX9 is not detected in young epidermis but is present in CK15^−^ basal keratinocytes of aged epidermis. **(E)** Percentage of KLF4^high^, GRHL3^high^, CK15^+^ and SOX9^+^ cells in young and aged epidermis (mean ± SEM; n = 3 per group). ***, P < 0.001; n.d., not detected; Student’s t-test. In (B–D), nuclei are counterstained with DAPI (blue), dashed lines mark the dermal–epidermal junction and the skin surface, and merged images are shown in the right panels.

In young healthy skin, KLF4 was detected in broadly KCs in the epidermis, most highly expressed in the top layer keratinocytes and weakly in CK15^+^ base layer stem cells on rete ridge (**Figure 5B**, top panel), an expression pattern consistent with single-cell RNAseq results. In aged skin, the KLF4^high^ population in the top layer was significantly reduced (**Figure 5B**, bottom panel; statistics in **Figure 5E**). High GRHL3 expression was observed in the top layers, and the GRHL3-high population was significantly reduced in aged skin (**Figure 5C**, bottom panel; **Figure 5E**).

In contrast, SOX9 was detected in basal-layer KCs in aged epidermis that had lost CK15, and was not detected in young skin (**Figure 5D&E**). This shift suggests the emergence of an alternative stem or progenitor-associated transcriptional state during epidermal aging, potentially reflecting compensatory or maladaptive responses to the loss of structural niche cues.

Notably, these observations of normal skin specimens closely parallel the transcriptional programs observed in our ex vivo geometric model, in which flat matrices resembling aged skin preferentially induced SOX9-associated programs, whereas concave rete ridge topographies supported differentiation-associated transcriptional networks including KLF4 and GRHL3. Taken together, these findings support a model in which epidermal architecture functions as a key biophysical regulator of stem cell fate, linking niche geometry to chromatin accessibility and downstream transcriptional programs that govern epidermal differentiation. Age-related flattening of rete ridges may therefore disrupt these geometry-dependent regulatory networks, contributing to impaired EpiSC function and altered differentiation programs in aging human skin.

## DISCUSSION

Epidermal stem cells recovered from curved, rete ridge–like collagen scaffolds showed an early molecular response characterized by increased expression of epidermal differentiation and keratinization genes, reduced expression of cell cycle genes, and altered chromatin accessibility. Previous studies have shown that physical cues influence epidermal differentiation (*18–23*). Our study extends this work by linking transcriptional and chromatin accessibility changes in a bioprinted model of rete ridge geometry and comparing those changes with age-associated differences in human epidermis. This study supports a role for three-dimensional niche geometry in regulating epidermal stem cell homeostasis. We showed that concave topography reprograms the transcriptional and chromatin states of EpiSCs, and that key features of this response are mirrored in young compared with aged human epidermis. These findings identify niche geometry as an instructive cue for epidermal stem cell fate and epidermal homeostasis, and provide mechanistic insight linking age-associated epidermal atrophy to impaired EpiSC function.

Skin ridges begin forming at around 12–13 weeks of fetal development, after epidermal stem and progenitor cells are already present (*44, 45*). Experimental and computational models indicate that stem cells preferentially localize to the bottoms of rete ridges through differential adhesion to the basement membrane (*46*), and external cues from the niche are crucial for normal stem cell proliferation and differentiation (*13, 47*). Our results show that concave curvature promotes differentiation-associated transcriptional programs while suppressing cell cycle–related networks. Integrated ATAC-seq and RNA-seq analysis identified candidate transcription factors important for curvature-induced gene pathway changes. These include the epidermis differentiation drivers KLF4 (*34, 38, 48*), GRHL3 (*43, 49*), MAFB (*49*), CEBPB (*50*) and GATA3 (*51*), which were upregulated in the curved model and reduced in aged skin (**Figures 4…5**). In contrast, several factors important for stem cell maintenance and proliferation were downregulated, including SOX9 (*36*), SOX6 (*52*), FOXP1 (*37*) and TEAD1 (*38*). Notably, SOX9 is a marker and pioneer master regulator of hair follicle stem cells, while adult epidermal stem cells were SOX9 negative (*53*). In vivo, the reduction or loss of CK15^+^ stem cells in aged skin was accompanied by the emergence of SOX9^+^ epidermal basal cells, which were not detected in young skin (**Figure 5**), suggesting a shift of pluripotent progenitor cell identity and behavior within the basal domain.

Together, these parallels suggest that aging does not simply suppress keratinocyte differentiation but erodes an ordered differentiation program that rete ridge niches support. Because rete ridges provide numerous concave niches and expand the basal surface area, their flattening would be expected to shift basal cells from a CK15^+^ to a SOX9^+^ state, and adversely affect differentiation programs and skin renewal.

The ability of the curved niche to promote EpiSC differentiation need not deplete the stem cell pool; rather, it may be a key step in maintaining epidermal homeostasis in young, healthy skin. Mesa and colleagues showed that differentiation of a single basal stem cell is followed by division of an adjacent neighbor with a lag time of 1-2 days (*54*). EpiSC self-renewal is therefore not a constitutive driver of homeostasis but is tuned to tissue demand and coupled to local differentiation and available space (*55*). In this framework, the curved niche that efficiently directs a subset of cells toward differentiation also sustains proliferation of neighboring stem cells and, consequently, tissue renewal. Conversely, flattening of rete ridges would weaken this coupling, leading to stalled turnover rather than preserved stemness.

What is the signal triggering differentiation of EpiSCs in the curved niche? Studies have identified YAP/TAZ as central mechanotransducers in the epidermis (*21*). Their nuclear activity depends on cell spreading and matrix anchorage, and they sustain EpiSC proliferation in part by repressing Notch-driven differentiation (*21*). In parallel, YAP1/TAZ–TEAD complexes maintain epidermal homeostasis by promoting basal cell proliferation while limiting the activity of KLF4 (*38*). In the curved niche, the downregulation of *TEAD1* and *TEAD4* and YAP/TAZ–TEAD targets *CTGF* and *CYR61*, the reduced expression of cell–matrix adhesion genes, and the induction of *KLF4* together with KLF4-associated increased chromatin accessibility at upregulated genes all fit this regulatory axis. We therefore propose that concave topography may signal through the YAP/TAZ–KLF4 regulatory axis to promote the exit of epidermal stem cells from the progenitor state and their entry into the differentiation program. Our model suggests translational opportunities in skin aging, wound repair and regenerative design. Preservation or restoration of epidermal niche geometry—through topographically engineered biomaterials or biomechanically informed skin substitutes (*56*)—may help sustain EpiSC function in aged or chronically injured skin. In wound healing, where epidermal architecture is transiently disrupted, re-establishing appropriate curvature cues could facilitate efficient re-epithelialization and differentiation. More broadly, incorporating defined geometric features into engineered skin models or grafts may improve their physiological relevance and regenerative capacity.

Several limitations should be considered. First, EpiSCs were seeded at a low density to avoid the formation of cell clumps within the curved niches, which limited the cell–cell interactions that are key to epidermal stratification and homeostasis. Second, our model lacks supporting cell types, including fibroblasts, immune cells, melanocytes and vasculature, as well as appendages and dynamic dermal–epidermal interactions, all of which shape the EpiSC niche in vivo. However, this simplicity is also a strength. Unlike organoids or monolayer cultures, our 3D-bioprinted platform allows precise control of niche geometry in human cells while maintaining physiologically relevant cell–matrix interactions. This allowed us to isolate curvature as a single variable and to attribute the observed transcriptional and chromatin changes to geometry itself, an approach that has been difficult to apply to human epidermis. Although the model does not reproduce the native tissue environment, single-cell transcriptomic analysis and Immunofluorescence staining of human skin specimens showed that key curvature-responsive programs are mirrored in young versus aged epidermis, supporting their physiological relevance. Moreover, aged human skin grafted onto young immunodeficient mice regains features of young epidermis, including epidermal thickness and proliferation (*1*), consistent with, but not proving, the idea that the niche environment is a key determinant of EpiSC function.

In summary, this work demonstrates that physical niche geometry is a fundamental determinant of epidermal stem cell identity and function. By integrating 3D bioprinting with multi-omic profiling, we reveal a link between tissue architecture, chromatin regulation and stem cell fate. These findings underscore the principle that form shapes the function of epidermal stem cells and suggest that restoration of structural complexity may offer new avenues to improve regenerative capacity in skin aging and wound healing.

## MATERIALS AND METHODS

### Human tissue samples

Formalin-fixed, paraffin-embedded samples of healthy, normal adult human skin were examined, including a young cohort (n = 6; aged 19-30 years) and an older cohort (n = 6; aged 79–92 years) (**Table S1**); three specimens per group were used for IF staining. Owing to the limited availability of biopsies from clinically normal human skin, these samples were obtained from histologically normal margins of excision specimens. The H&E-stained slides for each specimen were reviewed by dermatopathologists (C.G.L and G.F.M) to confirm the diagnoses and to specify the areas of normal skin for analysis. All human tissue samples were derived from the Pathology Archives of Brigham and Women’s Hospital with full institutional review board approval. Patient consent was not required because de-identified pathological specimens are considered discarded material by our institution, and these studies were therefore exempt.

### Histology and morphometry

Five μm sections were stained with H&E. Epidermal thickness was measured from the basement membrane to the base of the stratum corneum at the deepest point of rete ridges and at inter-ridge regions. The mean value per specimen was used for statistical analysis.

### Preparation of 3D-bioprinted scaffolds with a curved artificial “rete ridge” niche

Collagen scaffolds were fabricated using a microvalve-based robotic bioprinting platform (CLECELL, Seongnam-si, South Korea) as previously described (*57, 58*). The printer consists of four independently controlled liquid-dispensing channels, each operated by electromechanical valves mounted on a three-axis robotic stage. Liquid materials are dispensed by pneumatic pressure, with droplet size determined by parameters such as valve opening time and pressure. Bovine collagen (6 mg/mL; Sigma-Aldrich, St. Louis, MO) was loaded into a 10-cc syringe used as the printing cartridge. Pressures of 6–7 psi and a valve opening time of 500 μs were used, yielding droplets of approximately 50–100 nL with 500-μm spacing. Using a layer-by-layer fabrication method, a multilayered collagen scaffold was printed on 60-mm cell culture dishes (Techno Plastic Products, Switzerland). The dishes were uniformly coated with nebulized 1 M NaHCO_3_ before printing to increase collagen gelation and adhesion to the surface. Nebulized NaHCO_3_ was similarly applied after printing each layer, allowing approximately 1–2 min between layers for adequate gelation before deposition of the next layer. The final settings were a resolution of 650 μm, a valve opening time of 800 μs, and nebulizer settings of 70% power and 5-s treatment time. Both flat and curved scaffolds measured 18 mm × 18 mm. Curved scaffolds comprised 10 layers deposited in a zig-zag pattern, whereas flat scaffolds comprised 5 zig-zag layers overlaid by at least 5 “snail” rim layers, creating a gentle ridge around the edges to prevent spillover of cells during seeding (**Figure 1C**).

A resin stamp was designed and engineered using SLA 3D printing to imprint the collagen scaffolds with rete ridge–like topography. The stamp (18 mm × 18 mm) featured protrusions with a diameter of 100 μm, height of 800 μm and spacing of 600 μm (**Figure 1C**). These stamps were created and provided by Dr. Seung-Schik Yoo. Imprints were created by pressing the stamps (lubricated with sterile PBS) firmly into fully gelated collagen scaffolds for 5 s, then releasing manual pressure while keeping the stamp in place overnight until cell seeding. All scaffolds were incubated overnight at 37°C, 5% CO_2_, and stamps were gently removed before cell seeding.

### Isolation, seeding and harvesting of EpiSCs on flat and curved matrices

Primary human EpiSCs were isolated from neonatal foreskin using a one-step, serum-free method described previously(25). Cells were cultured in Keratinocyte Serum-Free Medium (SFM; Thermo Fisher Scientific, Waltham, MA) supplemented with human recombinant epidermal growth factor, bovine pituitary extract and penicillin–streptomycin (Thermo Fisher Scientific) and maintained for at least two passages. EpiSCs at passages 2–3 were seeded at 0.5 million cells per 100-mm Corning tissue culture–treated dish (Fisher Scientific) and maintained in supplemented Keratinocyte SFM. For seeding onto scaffolds, cells were detached with 1 mL 0.05% trypsin–EDTA (Gibco) for approximately 5–10 min and seeded dropwise onto the surface of curved and flat scaffolds at 100,000 cells per scaffold. After 1 h at 37°C, 5% CO_2_, scaffolds were covered with supplemented Keratinocyte SFM and incubated for 36 h before harvesting for RNA-seq and ATAC-seq. EpiSCs on flat scaffolds were harvested with a single treatment of 0.05% trypsin–EDTA, whereas those on curved scaffolds underwent one treatment to remove cells adhered to the intervening flat surfaces and a second to harvest the cell aggregates captured in the concavities (**Figure 1D**). Phase-contrast images were acquired at seeding, 36 h post-seeding and after harvesting.

### RNA-seq and data analysis

RNA-seq of EpiSCs harvested from flat or curved scaffolds (n = 3 per condition) was performed by GENEWIZ, LLC (South Plainfield, NJ). Samples were processed using the RNA-seq pipeline implemented in the bcbio-nextgen project. Raw reads were examined for quality using FastQC (*59*). Adapter sequences, other contaminant sequences such as polyA tails, and low-quality sequences with Phred quality scores below five were trimmed using Atropos (*60*). Trimmed reads were aligned to the UCSC human hg38 build, augmented with transcript information from Ensembl release GRCh38.p13, using HISAT2 (*61*). Alignments were checked for evenness of coverage, rRNA content, genomic context, complexity and other quality control parameters using FastQC (*59*), Qualimap (*62*), MultiQC and custom tools. Counts of reads aligning to known genes were generated using featureCounts (*63*), and differential expression was analyzed using DESeq2 (*64*). DEGs were defined by FDR < 0.05 and |fold change| ≥ 1.5. GO and KEGG enrichment of DEGs was examined using DAVID (*65*). GSEA (*66*) of genes ranked by expression change was performed using clusterProfiler (*67*). Expression of signature genes of basal (BAS-I to BAS-IV), spinous (SPN-I and SPN-II) and granular (GRN) keratinocytes defined by Wang et al. (*26*) was visualized as heatmaps of row-scaled expression.

### ATAC-seq and data analysis

ATAC-seq was performed by Active Motif, Inc. (Carlsbad, CA) on two independent preparations of EpiSCs harvested from curved scaffolds, flat scaffolds and conventional 2D culture. More than 30 million 40-bp paired-end reads were obtained per sample. Raw reads were quality-checked with FastQC (*59*), trimmed using Trim Galore (*68*) and aligned to the human reference genome (GRCh38/hg38) using BWA (*69*); alignments with mapping quality (MAPQ) > 20 were retained. ATAC-seq peaks were identified in each replicate and across biological replicates of each condition using MACS2 with default parameters for narrow peaks (*70*). To identify differential regions, peaks called in each replicate were merged, read counts at merged peaks were quantified using featureCounts (*63*) and normalized using the trimmed mean of M-values (TMM) method, and differential accessibility was assessed using a negative binomial generalized linear model with quasi-likelihood F-test in edgeR (*29, 30*). Regions were considered differentially accessible using two thresholds: FDR < 0.05 and |fold change| ≥ 1.5 (used in most analysis unless otherwise stated), or nominal P < 0.05 and |fold change| ≥ 1.5. Owing to the limited number of biological replicates, the nominal P < 0.05 threshold was used in examining the correlation of ATAC change and gene expression changes in **Figure 1I/J**. Genomic distribution of differential regions was annotated using ChIPseeker (*71*). Differential regions were assigned to genes with transcription start sites within ±5 kb, and the association between differential accessibility and differential expression was assessed by Fisher’s exact and chi-square tests (**Table S2**). Motif enrichment analysis was performed using HOMER (*72*), and protein–protein interaction networks of TFs corresponding to enriched motifs were generated using STRING (*73*). Consensus TF binding sites were obtained from ReMap 2022 (28). Published ChIP-seq datasets of H3K27ac (*74*), H3K4me1 and H3K4me3 (*75*) in neonatal foreskin keratinocytes, and H3K4me2 (*76*), KLF4 (*48*) and GRHL3 (*43*) in normal human epidermal keratinocytes (NHEK) were used to examine the epigenetic landscape of differential ATAC-seq regions. Signal heatmaps and profiles were generated using deepTools (*77*), and correlations between ChIP-seq signals and gene expression changes at differential ATAC-seq regions were assessed by Pearson’s correlation.

### Single-cell RNA-seq analysis

A published scRNA-seq dataset of human epidermis (*39*) (GEO GSE281449) was reanalyzed in R using Seurat 5.0.1. The dataset comprises three young donors (20, 21 and 24 years) and three old donors (61, 63 and 77 years). We removed predicted doublets (scDblFinder 1.16.0) and cells with fewer than 200 or more than 4,000 detected genes or more than 25% mitochondrial UMIs. We also removed genes detected in fewer than 3 cells, leaving 12,597 cells and 21,022 genes. Ambient RNA was corrected using (decontX, celda 1.28.0), and donor effects were corrected by Harmony 2.0.4. After log-normalization, cells were clustered (shared nearest-neighbor graph, Louvain algorithm, resolution 0.5) and identified with marker genes. To identify developmental stages, interfollicular keratinocytes (n=4,518) were re-normalized and clustered on the 2,500 most variable genes excluding 1,143 non-keratinocyte genes with ≥4-fold higher expression in non-keratinocytes than in keratinocytes. For differential expression analysis, raw UMI counts of each keratinocyte cluster were summed per donor to create pseudobulk counts. Donors with fewer than 10 cells in a cluster were excluded, as were genes with fewer than 10 counts (edgeR filterByExpr). Differentially expressed genes (DEGs) were identified using DESeq2 (*64*) and defined as nominal P < 0.05 and |fold change| ≥ 1.5 (young/old).

### Immunofluorescence staining

IF staining was performed on paraffin-embedded sections of formalin-fixed human skin and on EpiSCs in 2D culture. Five-μm paraffin sections were deparaffinized, rehydrated and heated in Target Retrieval Solution (Agilent Technologies) in a pressure cooker. Sections were blocked with 10% goat serum for 1 h and incubated overnight at 4°C with primary antibodies against CK15 (MA5-11344, Thermo Fisher Scientific), Ki-67 (PA1-21520, Thermo Fisher Scientific), FABP5 (33926, Cell Signaling Technology), KLF4 (HPA002926, Sigma-Aldrich), GRHL3 (HPA059960, Sigma-Aldrich) and SOX9 (ab185966, Abcam). Sections were washed, and incubated with secondary antibodies Alexa Fluor 488 goat anti-rabbit IgG (A11008, Invitrogen) and Alexa Fluor 647 goat anti-mouse IgG2a (A21241, Invitrogen). Slides were mounted with ProLong Glass Antifade Mountant with NucBlue (Invitrogen) and imaged using an EVOS FL Auto 2 microscope (P36981, Invitrogen). The percentage of KLF4^+^, GRHL3^+^, CK15^+^ and SOX9^+^ cells among epidermal keratinocytes was quantified in 3 fields per specimen. EpiSCs in 2D culture were fixed with 3% paraformaldehyde in PBS for 10 minutes, permeabilize with 0.2% triton x-100 in PBS for 20 minutes, and blocked and stained with antibodies as above.

### Statistical analysis

Data are presented as mean ± SEM. Comparisons between young and aged skin were performed using two-tailed Student’s t-test. Statistical methods for sequencing analyses are described above. P < 0.05 was considered significant.

