## Supplemental Materials for "Rete Ridge Topography Regulates Epidermal Stem Cell Differentiation Programs and Chromatin Accessibility"

**LIST OF SUPPLEMENTARY MATERIALS**

Fig S1. Measure of epidermal thickness of health skin from young and old donors.

Fig S2. Characterization of EpiSCs isolated from neonatal foreskin.

Fig S3. Pathway enrichment of differentially expressed genes in EpiSCs cultured in curved niche versus flat scaffold.

Fig S4. Chromatin accessibility in EpiSCs cultured in curved and flat gel matrix, or in standard 2D culture.

Fig S5. Protein–protein interaction network of transcription factors whose motifs are enriched at differential ATAC-seq regions in EpiSCs in the curved niche versus the flat matrix.

Fig S6. Epigenetic landscape at differential ATAC-seq regions defined at a less stringent threshold in EpiSCs in the curved niche compared with the flat matrix.

Fig S7. Single-cell transcriptomic analysis of keratinocyte subpopulations and transcription factor expression in young and aged human epidermis.

Table S1. Human young and old skin specimens.

Table S2. Association between differential chromatin accessibility and differential gene expression in EpiSCs in the curved niche versus flat gel matrix

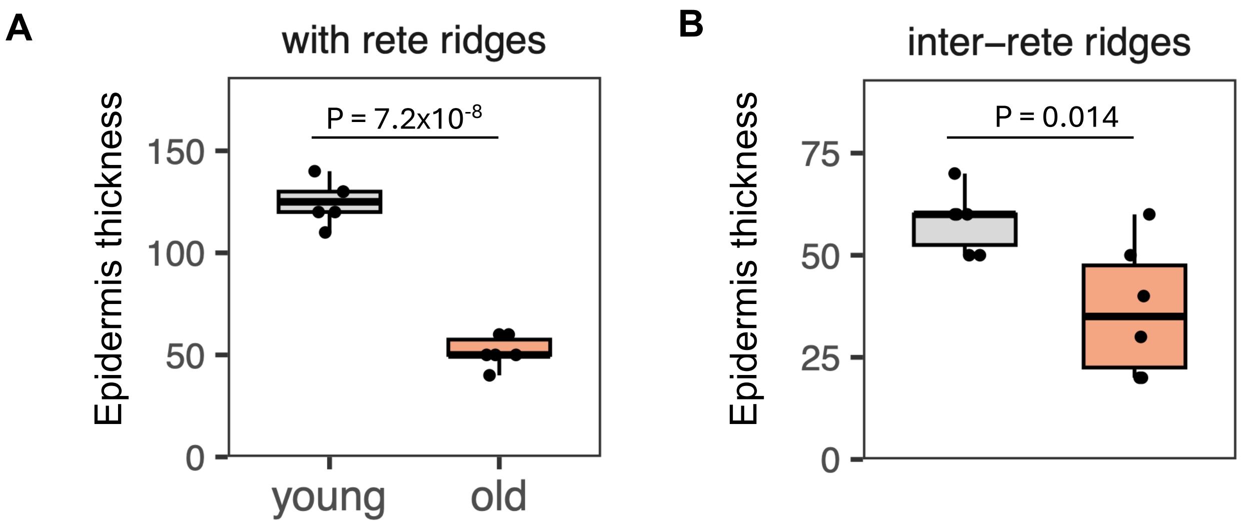


**Fig S1. Measure of epidermal thickness of health skin from young and old donors.** Bar graph plotted the average thickness of rete ridges (A) and inter-rete ridge regions (B) per donor. P, two-tail Student's t-test. N = 6 donors per group.

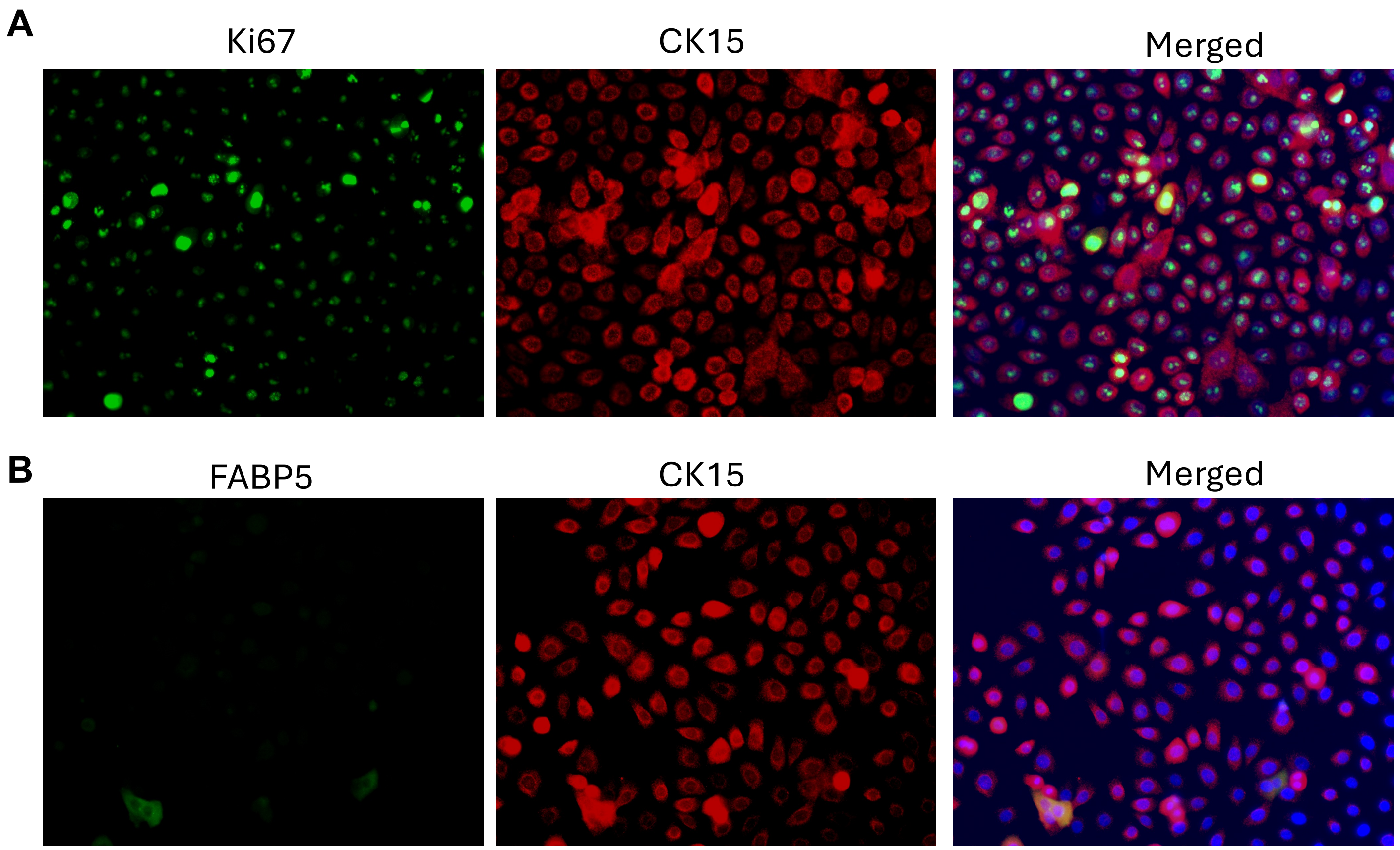


**Fig S2. Characterization of EpiSCs isolated from neonatal foreskin.**
Immunofluorescence staining of EpiSCs in conventional 2D culture for (B) Ki-67 (green) and CK15 (red), and (B) FABP5 (green) and CK15 (red). Ki-67 is present throughout the cell cycle (G1, S, G2 and M) and absent only in resting G0 cells. Most cells show Ki-67 nuclear staining. The punctate perinucleolar pattern marks cells in G1/early S phase, and the intense staining throughout the nucleus marks cells in G2/M. Both patterns were scored as Ki-67$^{+}$. Nuclei are counterstained with DAPI (blue).

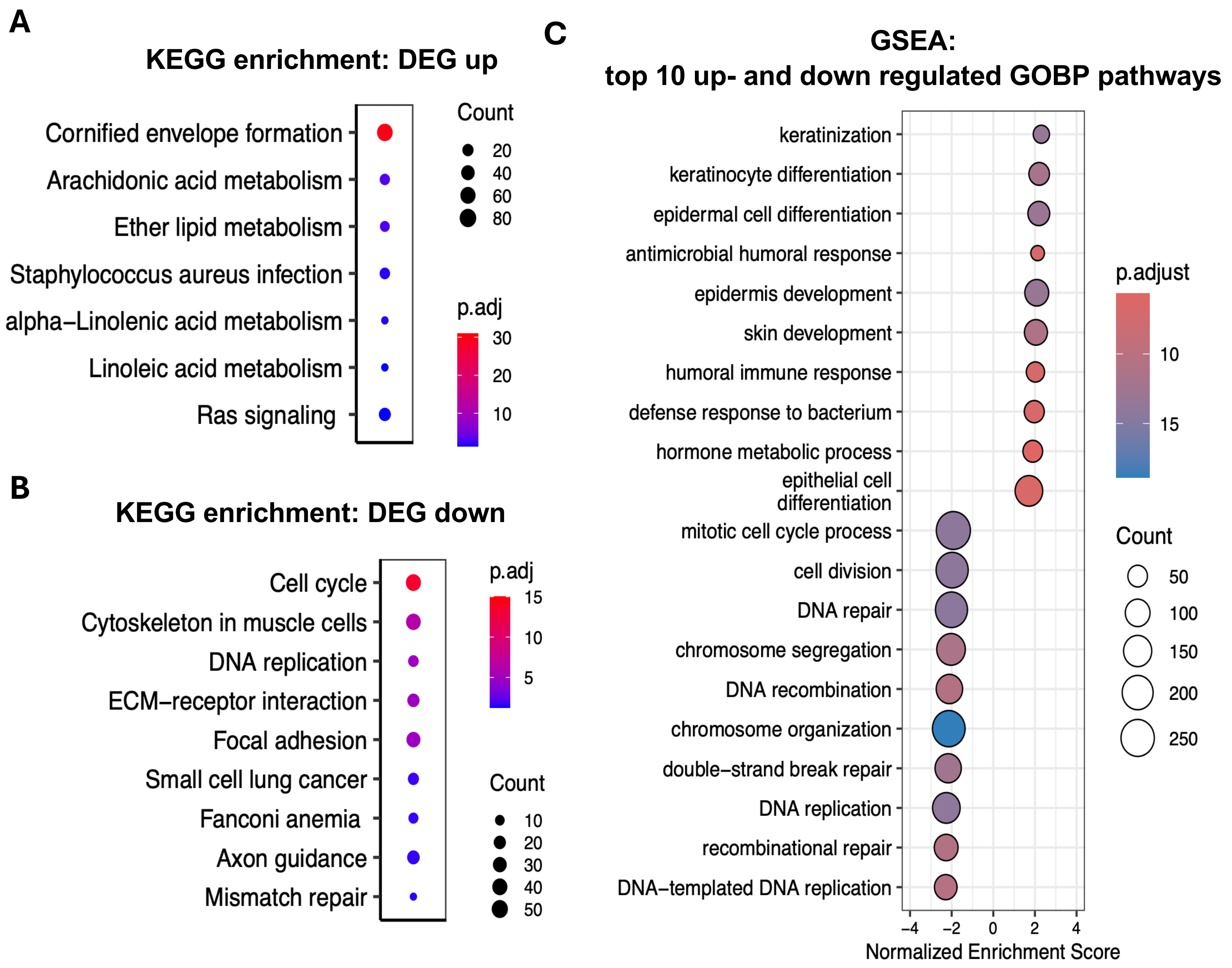


**Fig S3. Pathway enrichment of differentially expressed genes in EpiSCs cultured in curved niche versus flat scaffold.**

**A–B**. KEGG pathway enrichment of upregulated (A) and downregulated (B) DEGs. The top 10 pathways ranked by adjusted P value are shown.

**C**. GSEA of the top 10 upregulated and downregulated GOBP pathways.

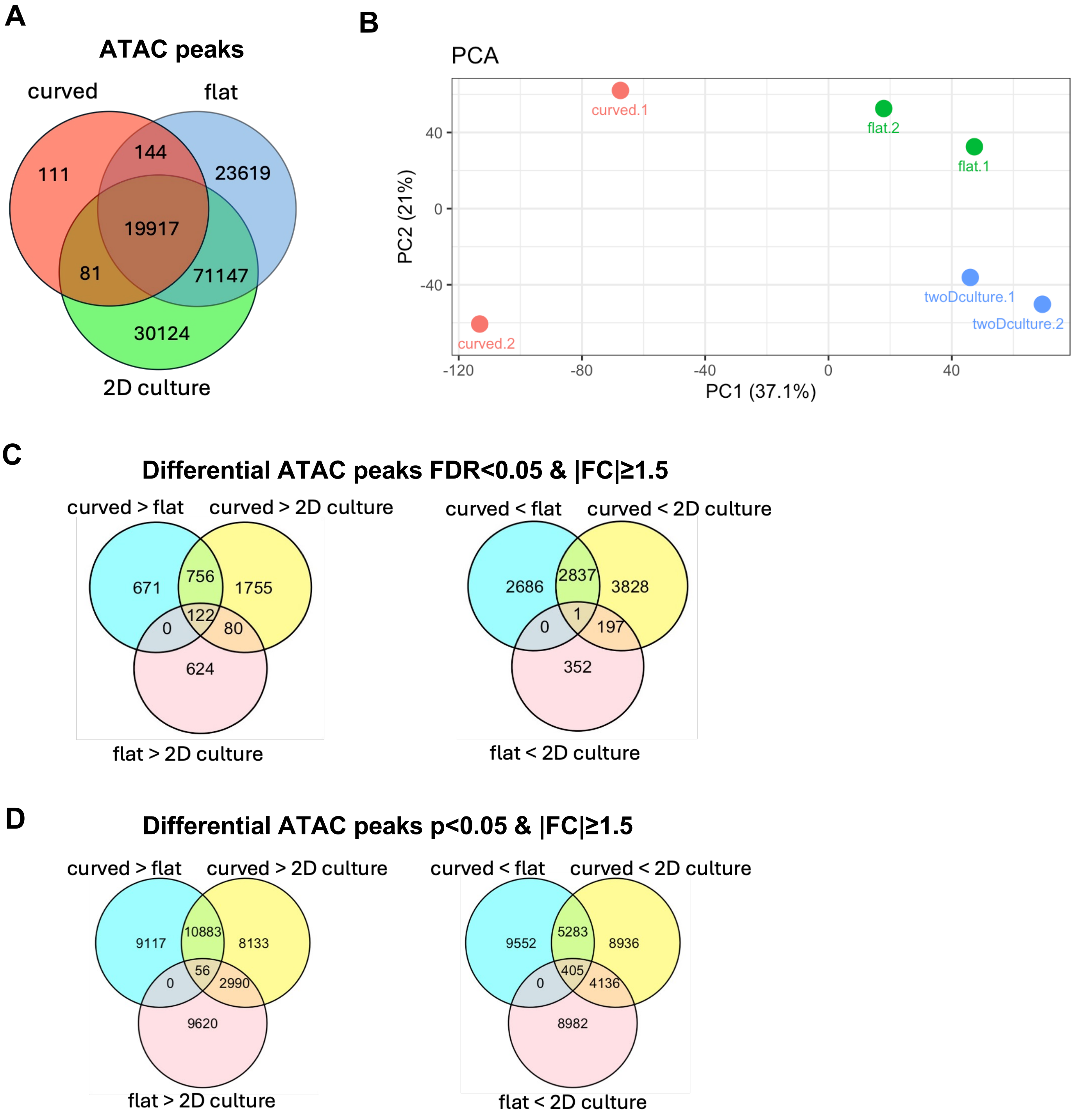


**Fig S4. Chromatin accessibility in EpiSCs cultured in curved and flat gel matrix, or in standard 2D culture.**

**A**. Venn diagram of ATAC-seq peaks identified in EpiSCs cultured in the curved niche, on a flat scaffold or in a standard culture dish (curved, flat and 2D culture, respectively).

**B**. Principal component analysis of read counts at merged ATAC-seq peaks (all peaks identified in either replicate of the three conditions).

**C-D**. Venn diagrams of differential ATAC-seq peaks between conditions, defined by FDR < 0.05 and |fold change| ≥ 1.5 (C) or by nominal P < 0.05 and |fold change| ≥ 1.5 (D).

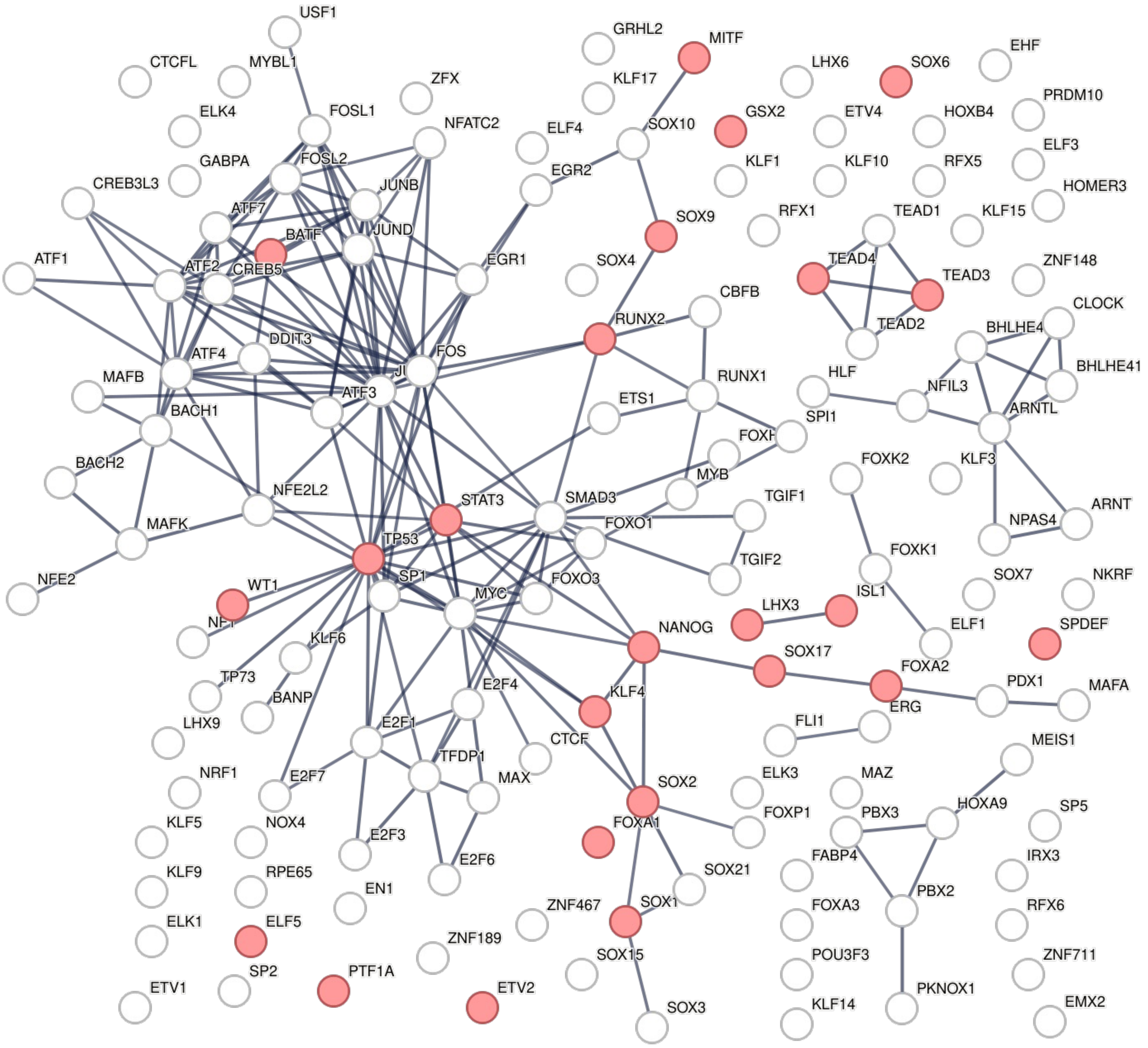


**Fig S5. Protein–protein interaction network of transcription factors whose motifs were enriched at differential ATAC-seq regions in EpiSCs in the curved niche versus the flat matrix.**

The STRING network shows transcription factors (TFs) whose motifs were enriched at diffPeaks (HOMER, adjusted P < 0.001). TFs annotated to the GO biological process cell fate commitment (GO:0045165) are shown in red.

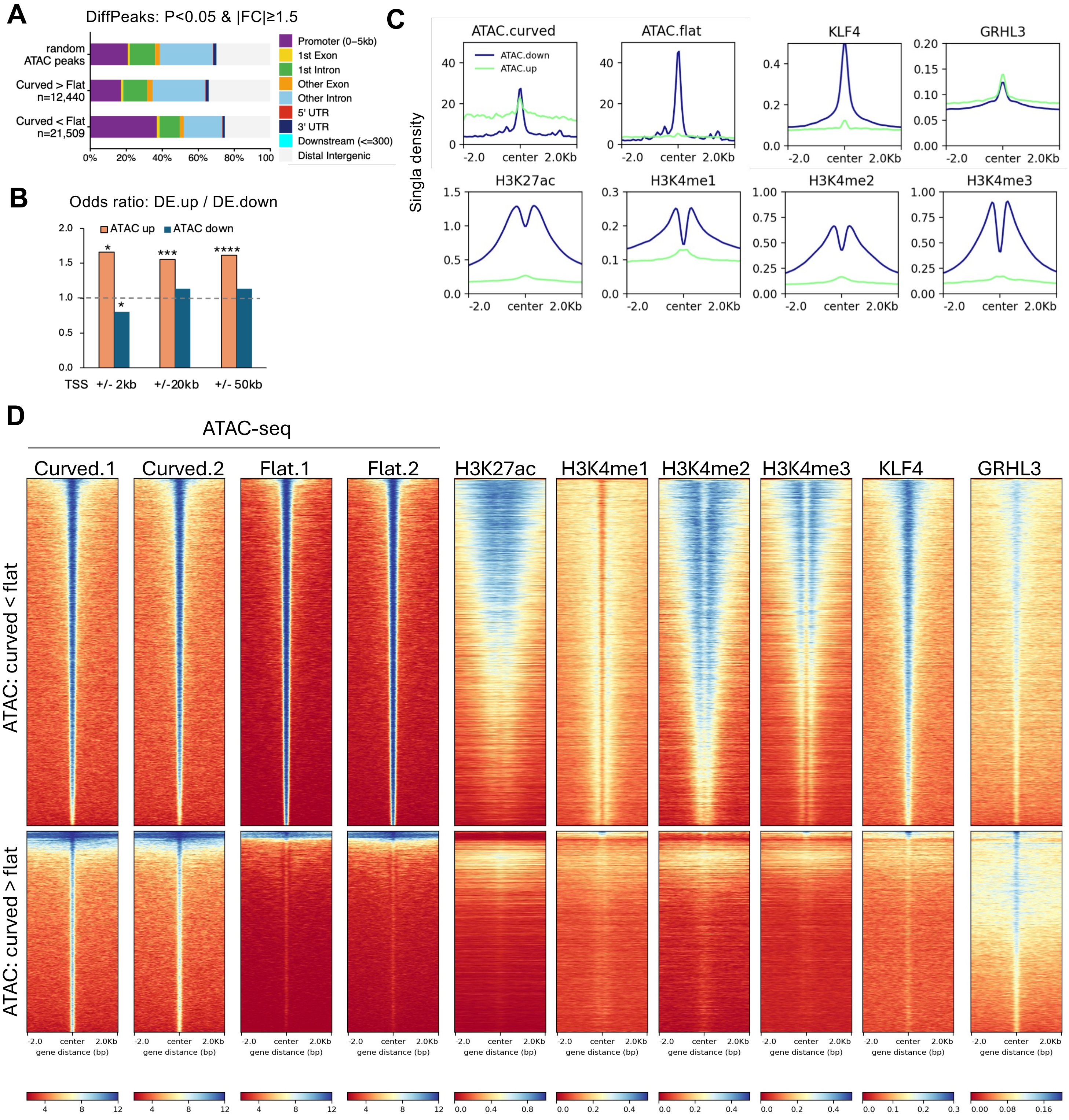


**Fig S6. Epigenetic landscape at differential ATAC-seq regions defined at a less stringent threshold in EpiSCs in the curved niche compared with the flat matrix.**

**A.** Genomic distribution of diffPeaks defined by nominal P < 0.05 and |FC| ≥ 1.5.

**B.** Bar graph of the odds ration of up- and down-regulated DEGs associated with diffPeaks with increased and decreased accessibility. *, chi-square p<0.05

**C.** Profile plots of ATAC-seq signals in EpiSCs on flat and curved matrices, and ChIP-seq signals of transcription factor (KLF4, GRHL3) and histone modifications in keratinocytes, centered on diffPeaks (±2 kb).

**D.** Heatmaps of the signals in (C).

Fig S6

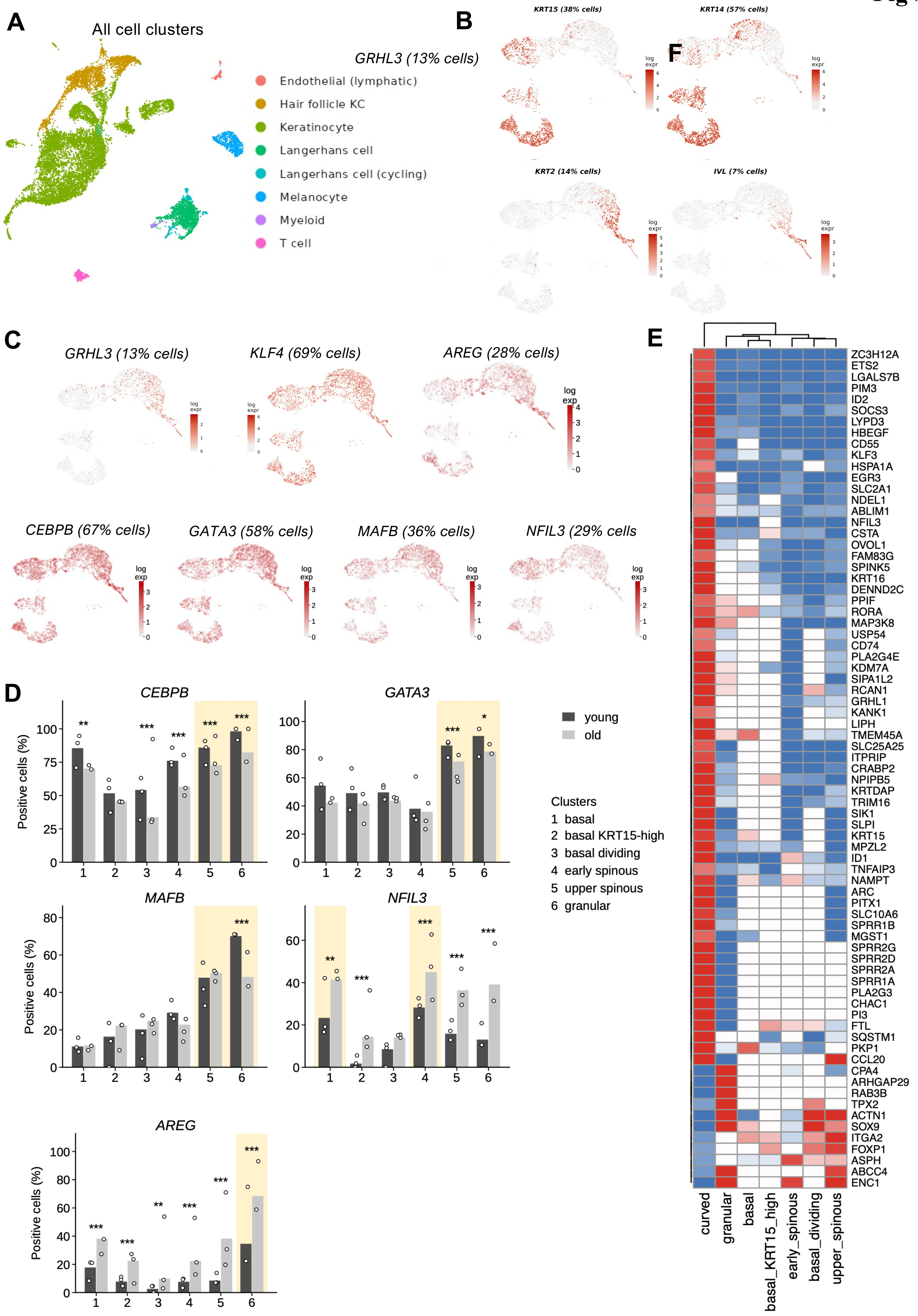
A
All cell clusters
GRHL3 (13% cells)
Endothelial (lymphatic)
Hair follicle KC
Keratinocyte
Langerhans cell
Langerhans cell (cycling)
Melanocyte
Myeloid
T cell
B
KRT15 (38% cells)
KRT14 (57% cells)
KRT2 (14% cells)
IVL (7% cells)
log expr
F
C
GRHL3 (13% cells)
KLF4 (69% cells)
AREG (28% cells)
CEBPB (67% cells)
GATA3 (58% cells)
MAFB (36% cells)
NFIL3 (29% cells
log exp
D
CEBPB
GATA3
MAFB
NFIL3
AREG
Positive cells (%)
young
old
Clusters
1 basal
2 basal KRT15-high
3 basal dividing
4 early spinous
5 upper spinous
6 granular
E
ZC3H12A
ETS2
LGALS7B
PIM3
ID2
SOCS3
LYPD3
HBEGF
CD55
KLF3
HSPA1A
EGR3
SLC2A1
NDEL1
ABLIM1
NFIL3
CSTA
OVOL1
FAM83G
SPINK5
KRT16
DENND2C
PPIF
RORA
MAP3K8
USP54
CD74
PLA2G4E
KDM7A
SIPA1L2
RCAN1
GRHL1
KANK1
LIPH
TMEM45A
SLC25A25
ITPRIP
CRABP2
NPIPB5
KRTDAP
TRIM16
SIK1
SLPI
KRT15
MPZL2
ID1
TNFAIP3
NAMPT
ARC
PITX1
SLC10A6
SPRR1B
MGST1
SPRR2G
SPRR2D
SPRR2A
SPRR1A
PLA2G3
CHAC1
PI3
FTL
SQSTM1
PKP1
CCL20
CPA4
ARHGAP29
RAB3B
TPX2
ACTN1
SOX9
ITGA2
FOXP1
ASPH
ABCC4
ENC1
curved
granular
basal
basal_KRT15_high
early_spinous
basal_dividing
upper_spinous

**Fig S7. Single-cell transcriptomic analysis of keratinocyte subpopulations and transcription factor expression in young and aged human epidermis.**
**A.** UMAP of all cell types in young and aged skin, colored by cluster. Keratinocytes were extracted and re-clustered to define the subclusters shown in Figure 4A.
**B.** UMAP feature plots of established keratinocyte differentiation markers.
**C.** UMAP feature plots of transcription factors and AREG in keratinocyte subclusters.
**D.** Percentage of cells expressing each gene in keratinocyte subclusters of young and aged skin. Shading indicates the subcluster(s) with the highest expression. Fisher's exact test with Benjamini–Hochberg correction; *FDR < 0.05, **FDR < 0.01, ***FDR < 0.001.
**E.** Heatmap of log2 fold changes of genes that changed in opposite directions in young (vs. aged) skin keratinocytes and in EpiSCs in the curved niche (vs. flat matrix). Values are capped at ±1, and changes smaller than 10% ($|log2FC| < 0.14$) are shown in white.

**Table S1. Human young and old skin specimens.**

| Group | Age | Sex | Site |
|---|---|---|---|
| Young | 19 | F | Arm |
| | 20 | F | Arm |
| | 26 | M | Neck |
| | 30 | M | Shoulder |
| | 21 | F | Abdomen |
| | 22 | M | Chest |
| Old | 83 | M | Face |
| | 79 | M | Thigh |
| | 92 | F | Shoulder |
| | 82 | M | Chest |
| | 86 | F | Back |
| | 83 | M | Face |

**Table S2. Association between differential chromatin accessibility and differential gene expression in EpiSCs in the curved niche versus flat gel matrix**

| diffPeak | peak set | distance | associated genes | associated DEG.up | associated DEG.down | Normalized odds ratio # | p value* |
|---|---|---|---|---|---|---|---|
| p<0.05 & \|FC\|>=1.5 | ATAC up | TSS ± 2kb | 689 | 52 | 29 | **1.66** | **0.031** |
| | ATAC down | | 6180 | 377 | 394 | **0.8** | **0.012** |
| | ATAC up | TSS ± 50kb | 7057 | 558 | 389 | **1.62** | **$6.8\times10^{-8}$** |
| | ATAC down | | 13015 | 864 | 761 | 1.14 | 0.24 |
| FDR<0.05 & \|FC\|>=1.5 | ATAC up | TSS ± 2kb | 16 | 1 | 0 | -- | 1 |
| | ATAC down | | 2542 | 152 | 161 | 0.83 | 0.14 |
| | ATAC up | TSS ± 50kb | 254 | 32 | 12 | **2.46** | **0.0087** |
| | ATAC down | | 6947 | 505 | 386 | 1.35 | 0.00086 |

#, Odds ratio = (associated DEG.up / associated DEG.down) / (non-associated DEG.up / non-associated DEG.down

*, Two-sided Fisher's exact test.

Blue, concordant changes in chromatin accessibility and gene expression (increased accessibility with upregulation, or decreased accessibility with downregulation).
